# A Global Genomic Resource for Outcrossing *Arabidopsis lyrata* and *Arabidopsis arenosa*

**DOI:** 10.64898/2026.03.02.709016

**Authors:** Anna Glushkevich, Laura Steinmann, Nikita Tikhomirov, Jakub Vlček, Yu Cheng, Jana Flury, Uliana Kolesnikova, Miloš Duchoslav, Jörn Gerchen, Gabriela Šrámková, Roman Ufimov, Sonia Celestini, Saurabh Pophaly, Magdalena Bohutínská, Veronika Lipánová, Levi Yant, Tiina Mattila, Roswitha Schmickl, Alison Scott, Filip Kolář, Polina Yu. Novikova

## Abstract

Genetic studies leveraging natural variation in *Arabidopsis* species have improved our understanding of evolutionary genetic processes underlying ecologically important and adaptive traits. Integrating the thorough functional knowledge accumulated in *A. thaliana* with the extensive natural variation in outcrossing *Arabidopsis* species is a powerful approach to study the basis of adaptation in natural evolutionary and ecological contexts. Here we present an integrated genomics database of sequenced genomes from several studies in *A. lyrata* (1018 genomes in total) and *A. arenosa* (736 genomes in total), spanning the geographic ranges of these two ploidy-variable, outcrossing taxa. We provide a searchable genome browser with population data mapped to respective reference genomes, an interactive geographic map of population structure clusters, and an efficient way to subsample the full dataset of genetic variation, available at arabidopsislyrata.org. To demonstrate its utility, we perform a genome-wide association study on a latitudinal cline of *A. lyrata* and find strong associations of several loci with latitude, including variants in key regulators of photoperiodic growth. This resource provides access to genetic diversity data in a single repository, enabling further studies of comparative genetics and local adaptation, as well as of individual genes of interest.

## Introduction

Understanding the genetic basis of plant adaptation to external factors (biotic or abiotic environments) and internal changes (whole genome duplications, mating system shifts), requires comprehensive, population-scale data on natural genetic variation sampled across the species’ entire geographic range. The sequenced collection of natural inbred lines of *Arabidopsis thaliana* proved to be a powerful tool for association studies between known genotypes and different physiological phenotypes or environments, measured in different laboratories (Atwell et al. 2010; Hancock et al. 2011; Long et al. 2013; The 1001 Genomes Consortium 2016). The relatives species of *A. thaliana* are outcrossing and require re-sequencing for every genotype-phenotype association study, but advances in high-throughput sequencing technologies minimized these disadvantages. Moreover, outcrossing *Arabidopsis* species are attractive as their higher genetic variation and effective recombination provide better resolution for genomic studies (Yant and Bomblies 2016) even with lower sample sizes compared to *A. thaliana*. While every model has pros and cons, multiple species offer an additional dimension and allow for comparative genomics of local adaptation (Whiting et al. 2024) among different lineages.

The *Arabidopsis* genus has a reticulated structure with many shared alleles across species due to their ancient and ongoing introgression (Novikova et al. 2016). The evolutionary history of *Arabidopsis* is about six million years old and started in Europe with multiple independent colonizations of Eastern parts of Eurasia, Africa and North America (Schmickl et al. 2010; Durvasula et al. 2017; Scott et al. 2025; Suda et al. 2025). *A. thaliana*, *A. lyrata*, *A. arenosa* and *A. halleri* are the most widespread *Arabidopsis* species with partially overlapping geographical distributions (Hoffmann 2005; Hohmann et al. 2014), while *A. cebennensis*, *A. pedemontana* and *A. croatica* are limited in their geography and genetic diversity (Novikova et al. 2016). Natural variation in outcrossing *Arabidopsis* species encompasses many additional traits and environmental gradients that are less pronounced or completely lacking in natural *A. thaliana* populations, such as ploidy variation (*A. arenosa, A. lyrata*) (Kolář, Lučanová, et al. 2016; Scott et al. 2025), mating system shifts (*A. lyrata) (Mable et al. 2005; Foxe et al. 2010; Griffin and Willi 2014; Kolesnikova et al. 2023; Li et al. 2023; Lucek et al. 2025)*, adaptation to toxic soils (*A. arenosa, A. lyrata, A. halleri) (Hanikenne et al. 2008; Turner et al. 2010; Arnold et al. 2016; Preite et al. 2019; Bohutínská et al. 2021),* or high-alpine temperate habitats (*A. arenosa) (Bohutínská et al. 2021; Wos et al. 2022)*. Moreover, genomics studies of adaptation in *A. lyrata* are further facilitated by extensive knowledge on local adaptation (Leinonen et al. 2009; Leinonen et al. 2011; Vergeer and Kunin 2011; Hämälä et al. 2018), and population differentiation in ecologically important traits (Kivimäki et al. 2007; Davey et al. 2008; Kuittinen et al. 2008; Huttunen et al. 2010; Sletvold et al. 2010; Vergeer and Kunin 2011; Sletvold and Ågren 2012; Quilot-Turion et al. 2013; Remington et al. 2013; Vergeer and Kunin 2013; Paccard et al. 2014; Hamala et al. 2017; Kemi et al. 2019; Heblack et al. 2024). Comparing the genetics underlying local adaptation across several species gives an idea of the extent to which such adaptation, although usually polygenic, can be repeated (Bohutínská and Peichel 2024; Celestini et al. 2025) and whether repeated solutions evolve independently by distinct mutations hitting the same genes, or by reusing the same allele shared as standing variation, or via introgression (Bohutínská and Peichel 2024; Celestini et al. 2025; Lee and Coop 2017).

Usually, even when raw short-read data from relevant species are available, reusing the data for comparative studies requires vast computational resources for storage and computing time: downloading the raw data, assembling and databasing their metadata, mapping, variant calling, and combining. Here, we provide full access to the up-to-date genotype matrices of available *A. lyrata* and *A. arenosa*, mapped to their respective reference genomes, as well as basic metadata on the original population’s sampling locations. We also provide easy interactive tools for visual inspection of the natural variation at the loci of interest and a quick way to cut and slice the matrix at the locus of interest for further analysis.

## Results and Discussion

### Functionality of the resource: browse, select and download

We obtained two separate genotype matrices: *A. lyrata* and *A. arenosa*. For the *A. lyrata* dataset, we mapped 779 published (Novikova et al. 2016; Mattila et al. 2017; Guggisberg et al. 2018; Hämälä et al. 2018; Hämälä and Savolainen 2019; Marburger et al. 2019; Takou et al. 2021; Willi et al. 2022; Kolesnikova et al. 2023; Bohutínská et al. 2024; Scott et al. 2025) and 131 newly sequenced *A. lyrata* samples, as well as 108 samples of closely related species (The 1001 Genomes Consortium 2016; Paape et al. 2018), to the *A. lyrata* NT1 reference genome of a self-compatible Siberian lineage (Kolesnikova et al. 2023). We chose the NT1 assembly over the previously used North American MN47 assembly, because the NT1 genome more closely resembles ancestral structure, while MN47 has large private structural variants (Hu et al. 2011; Kolesnikova et al. 2023). The closely related species include outgroups (*Capsella rubella*) and representatives of each species within the *Arabidopsis* genus in order to easily classify variants into ancestral and derived. We also included allotetraploid *A. kamchatica* to ensure differentiation of autotetraploid *A. lyrata* from *A. kamchatica*. The relatedness between the samples is represented by the distance matrix heatmap (fig. 1), the neighbor-joining network (supplementary fig. S1a) and PCA analysis (supplementary fig. S2). We also added methylation data and expression data (Hämälä et al. 2022; Kolesnikova et al. 2023), as well as long read RNAseq for the reference accession, as separate tracks to the browser for a possibility of a comprehensive analysis.

**Fig. 1.**
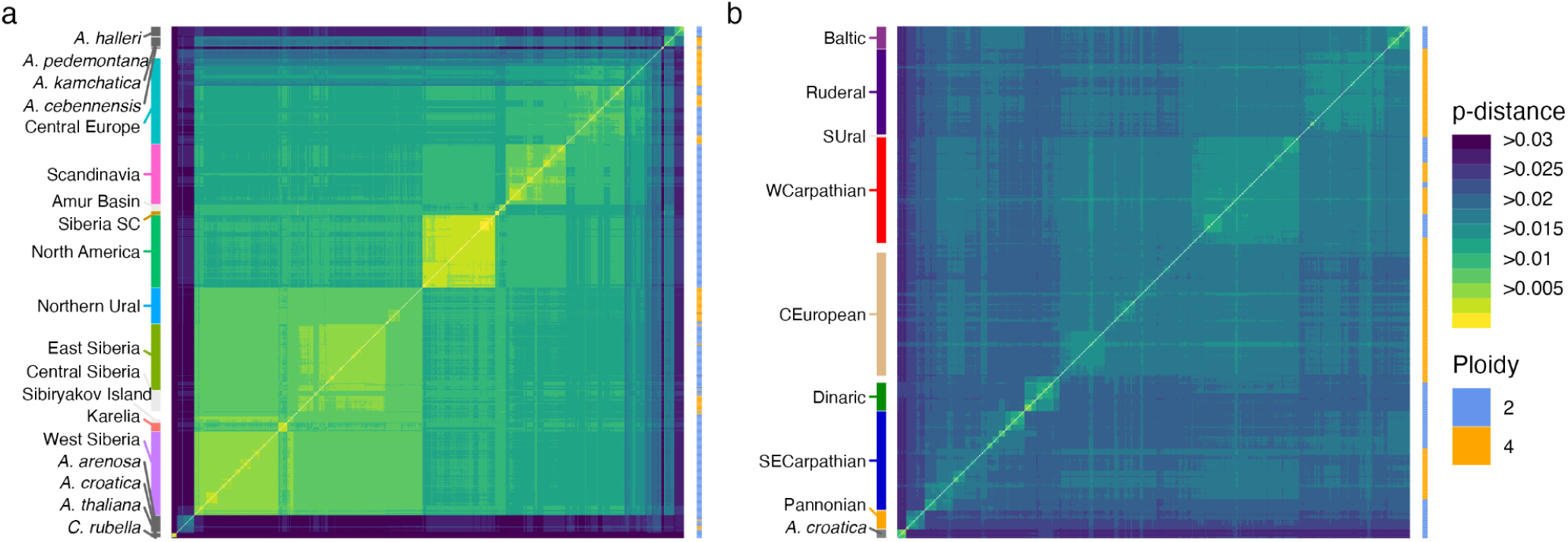
Genetic distance heatmaps of *Arabidopsis* species together with outgroups: (a) 1018 samples of *A. lyrata* and other species; (b) 723 *A. arenosa* and 13 *A. croatica* samples. The analysis is based on VCF2Dis p-distances calculated over synonymous four-fold degenerate sites. The blue/orange colour on the right of each panel corresponds to ploidy (2x/4x), the colours on the left denote the major lineages following cluster assignments in fig. 3. “Siberia SC” marks the self-compatible Siberian *A. lyrata* lineage.

The *A. arenosa* dataset comprises 736 individuals, including 13 individuals belonging to sister species *A. croatica.* The total dataset combines published (Hollister et al. 2012; Yant et al. 2013; Arnold et al. 2016; Novikova et al. 2016; Monnahan et al. 2019; Preite et al. 2019; Bohutínská et al. 2021; Konečná et al. 2021; Weitz et al. 2021; Bohutínská et al. 2024) and newly sequenced samples (fig. 1b). All samples were mapped to the reference genome of diploid *A. arenosa* from Pusté Pole, Western Carpathians, Slovakia (Bramsiepe et al. 2023). The relatedness between the samples is represented by distance matrix heatmap (fig. 1), the neighbor joining network (supplementary fig. S1b) and PCA analysis (supplementary fig. S3).

The relatedness between *Arabidopsis* species is highly reticulated, which is also confirmed by our species tree-gene tree reconciliation analysis based on 903 single-copy genes alignments (see Methods, supplementary fig. S4). Based on this approach, *A. arenosa* and *A. croatica* form a clade, which is sister to a clade of *A. cebennensis*, *A. pedemontana*, *A. halleri*, and *A. lyrata*.

The matrices include variant and non-variant sites called and filtered in a consistent way. Filtered matrices are searchable by TAIR gene nomenclature in the genome browser (arabidopsislyrata.org, Genome Browser) implemented with JBrowse 2 (Diesh et al. 2023) (fig. 2). The raw and prefiltered variant call format (VCF) files are also available for a download entirely or by selecting certain individuals and a certain locus coordinates (e.g., scaffold_1:24188299-24194609) for a download (arabidopsislyrata.org, VCF Selector).

**Fig. 2.**
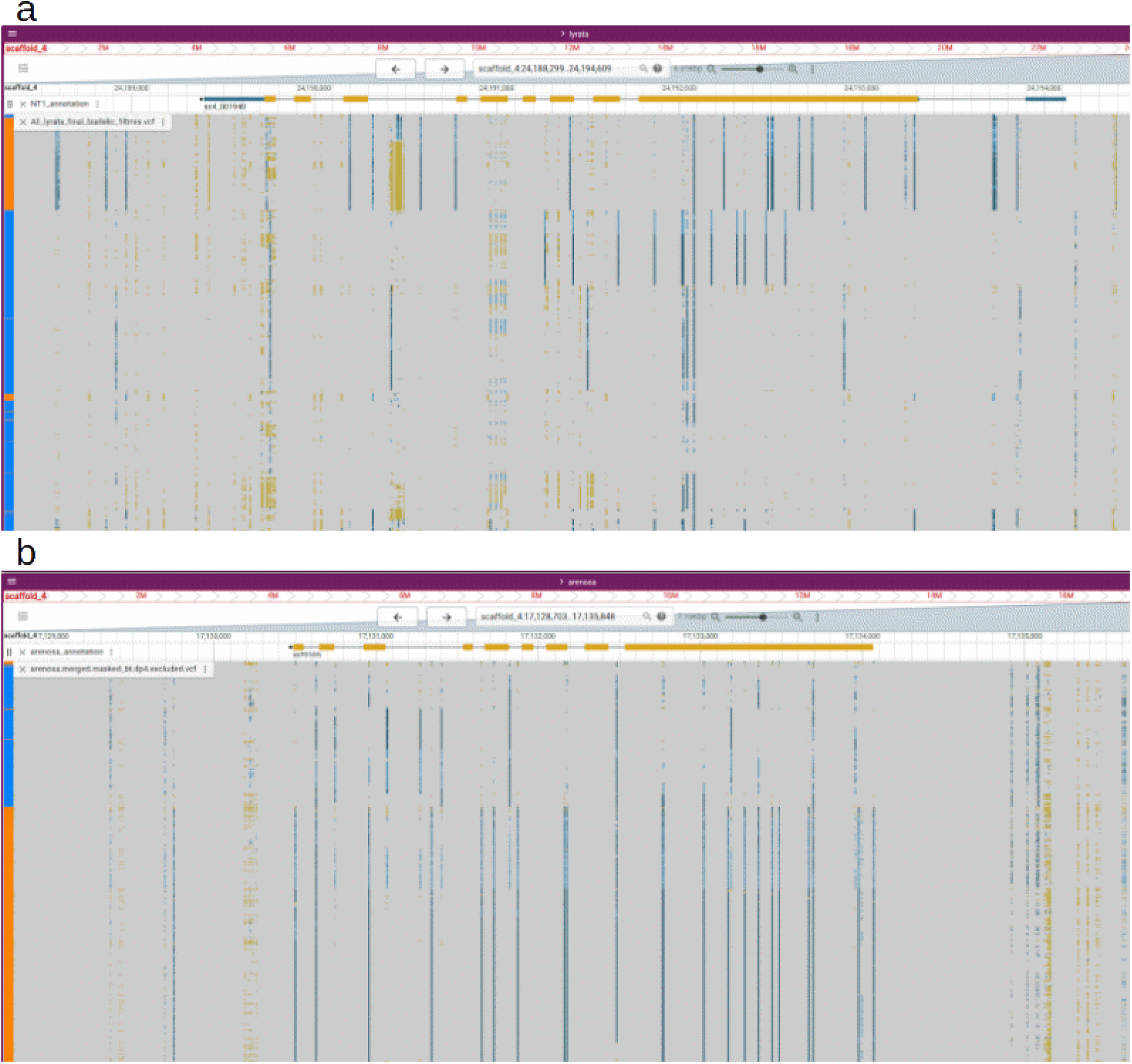
Genome browser screenshots of *A. lyrata* (a) and *A. arenosa* (b) matrix at ASY3 gene, involved in homologous chromosome pairing in meiosis and highly differentiated between diploid and autotetraploid populations (Yant et al. 2013; Morgan et al. 2020; Scott et al. 2025). The left panel represents individual ploidy: 2x (blue) and 4x (orange). Clustering of the genotypes matches ploidy, which is expected for this gene as it is known to have a tetraploid-specific haplotype (Yant et al. 2013; Arnold et al. 2016; Marburger et al. 2019; Monnahan et al. 2019; Scott et al. 2025).

To connect a particular genotype with both geographic origin of the individual and ancestry assignment, we provide interactive maps for both datasets with markers for each individual corresponding to assignments by designated genetic clustering methods (see Methods) (arabidopsislyrata.org Admixture Map) (fig. 3). In the default “clusters” representation, contribution of all ancestry groups is shown for each population sample, with detailed information on ancestry groups available on hover. In “groups” representation (toggled in the top right of the page), each population sample is assigned to one cluster with dominant ancestry contribution, with further metadata available on click.

**Fig. 3.**
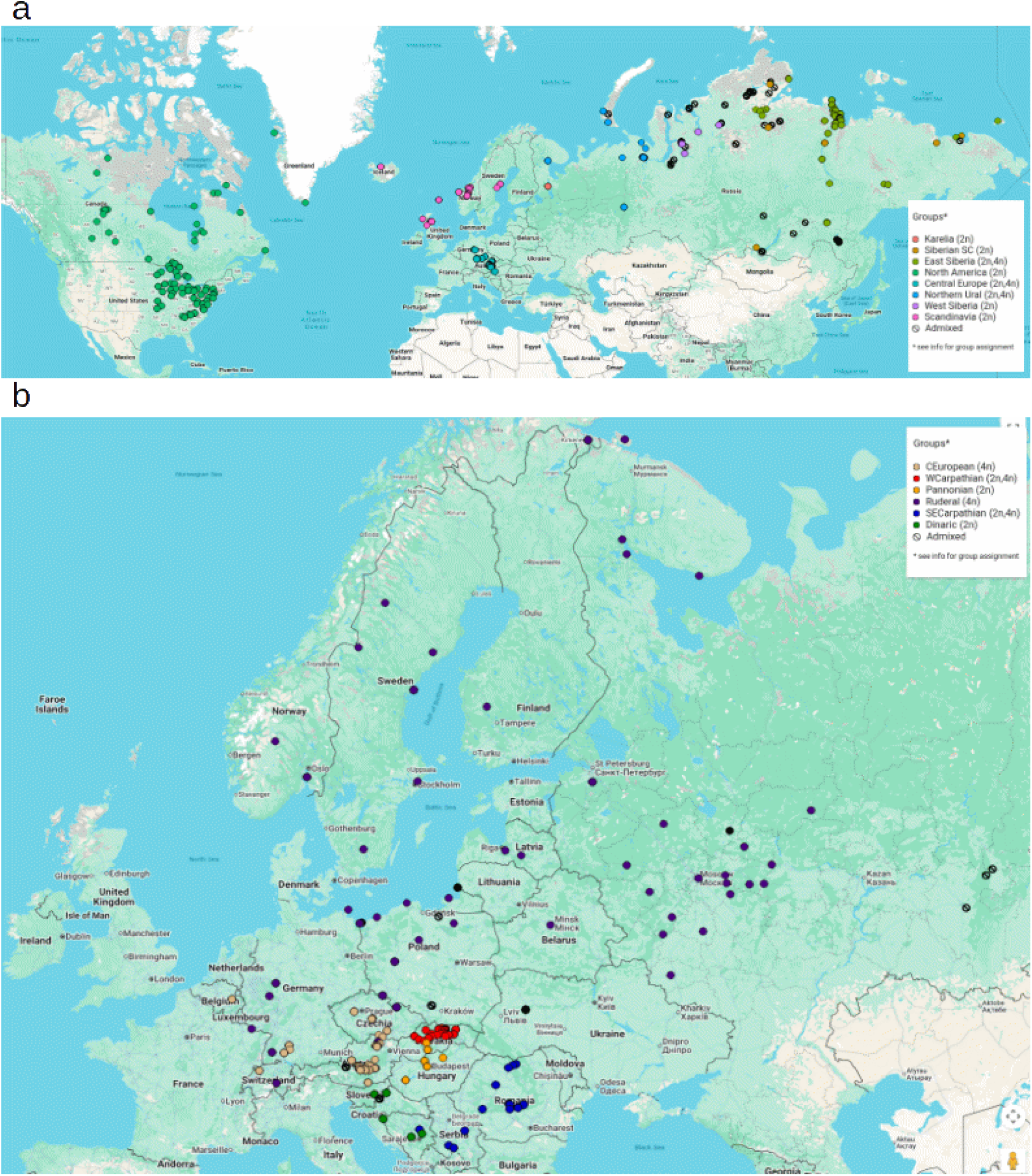
Distribution map of populations with summarised proportions of individual assignments to genetic groups for *A. lyrata* (eight genetic groups, k=8) (a) and *A. arenosa* (six genetic groups, k = 6) (b).

### Population structure and evolutionary history

For better orientation in the dataset we provide information on population structure within each species. *A. lyrata* and *A. arenosa* both have higher diversity than the cosmopolitan and mostly commensal selfer *A. thaliana* (The 1001 Genomes Consortium 2016; Hsu et al. 2019). Overall, genetic distance (fig. 1, supplementary fig. S1) and PCA analyses (supplementary fig. S2, S3) based on 4-fold degenerate sites split *A. lyrata* (fig. 3a) and *A. arenosa* (fig. 3b) into several distinct clusters reflecting geographical distribution and evolutionary history of each species (Monnahan et al. 2019; Scott et al. 2025).

The major splits in *A. lyrata* (K = 3) separate European, Asian (including NE European, i.e. Karelian) and Northern American populations, regardless of ploidy and/or traditional taxonomic delimitations (see Methods and supplementary fig. S2, S5). Under finer scale clustering (K = 8) presented in the interactive map (fig. 3, http://arabidopsislyrata.org, Admixture Map), additional geographical sub-clusters appear (Scandinavian and several Siberian). Both in Siberia and in Central Europe, tetraploids always belong to a mixed-ploidy cluster that also encompasses some diploid populations (fig. 3a). This is consistent with multiple whole-genome duplication events within *A. lyrata* that resulted in several stable autopolyploid lineages in Central Europe (Schmickl and Koch 2011; Bohutínská et al. 2024), Northern Ural and Central Siberia (Scott et al. 2025). *A. lyrata* is primarily an obligate outcrosser, including the autotetraploid lineages (Scott et al. 2025; Vekemans et al. 2025), with at least two independent transitions to self-compatibility (SC) in diploids: one in Siberia ∼90Kya (Kolesnikova et al. 2023), separate SC cluster (fig. 1a, fig. 3a), and one or more in North America likely following the last glacial maximum ∼10 kya (within the North American cluster fig. 2a, fig. 3a) (Mable et al. 2005; Foxe et al. 2010; Willi and Määttänen 2010; Griffin and Willi 2014). In addition, the recently diverged northeastern North American taxon *A. arenicola* (Willi et al. 2022) falls within the Northern American *A. lyrata* cluster (Supplementary fig. 5). Overall, the complex intraspecific structure of *A. lyrata* (fig. 1a, fig. 3a) reflects its colonisation history, spreading from Europe across Eurasia and Beringia all the way to North America (Ross-Ibarra et al. 2008; Schmickl et al. 2010; Pyhajarvi et al. 2012; Scott et al. 2025).

*A. arenosa* splits into six major lineages presented in the interactive map: two early-diverging diploid-only groups occupying Pannonian basin (Central Europe) and Dinaric mountains (Southeastern Europe), one exclusively tetraploid lineage (Central European) and three mixed-ploidy lineages (fig. 1b, fig. 3b, supplementary fig. 6). These results correspond with previous studies that identified origin of a tetraploid cytotype in Western Carpathians∼20-31k generations ago, that was followed by massive interploidy introgression in three regions that resulted in the current reticulate clustering by both geography and ploidy (Arnold et al. 2015; Monnahan et al. 2019). Indeed, one mixed-ploidy cluster represents the ancestral area of the autotetraploid lineage, where it still co-occurs and hybridizes with its diploid progenitor lineage (Western Carpathians (Arnold et al. 2015)). The other two mixed-ploidy lineages come from areas of secondary admixture where the expanding autotetraploid cytotype encountered distinct diploid lineages and crossed with them (Monnahan et al. 2019): Ruderal-Baltic lineage (mainly human-disturbed ruderal stands across Central and Northern Europe) and Southeastern Carpathian lineage restricted to SE Europe. While diploid individuals are generally well-delimited, showing assignment to a single cluster, a majority of the tetraploid individuals show high admixture (fig. 3b, supplementary fig. 2) indicating asymmetric interploidy gene flow mainly from diploids to tetraploids.

Finally, tetraploid *A. lyrata* and *A. arenosa* are known to hybridize and introgress in Central Europe (Schmickl and Koch 2011; Arnold et al. 2016; Novikova et al. 2016; Marburger et al. 2019), which is also reflected on the close positioning of Central European tetraploid *A. lyrata* and *A. arenosa* lineages on the network (supplementary fig. S1a).

### Contrasting patterns of genetic diversity in *A. lyrata* and *A. arenosa*

We assembled the most comprehensive whole-genome datasets to date for *A. lyrata* and *A. arenosa*. To characterise diversity, we calculated synonymous nucleotide diversity per species and population (fig. 4). Overall genetic diversity is higher in *A. arenosa* than in *A. lyrata* irrespective of ploidy, consistent with previous estimates (Novikova et al. 2016).

**Fig. 4.**
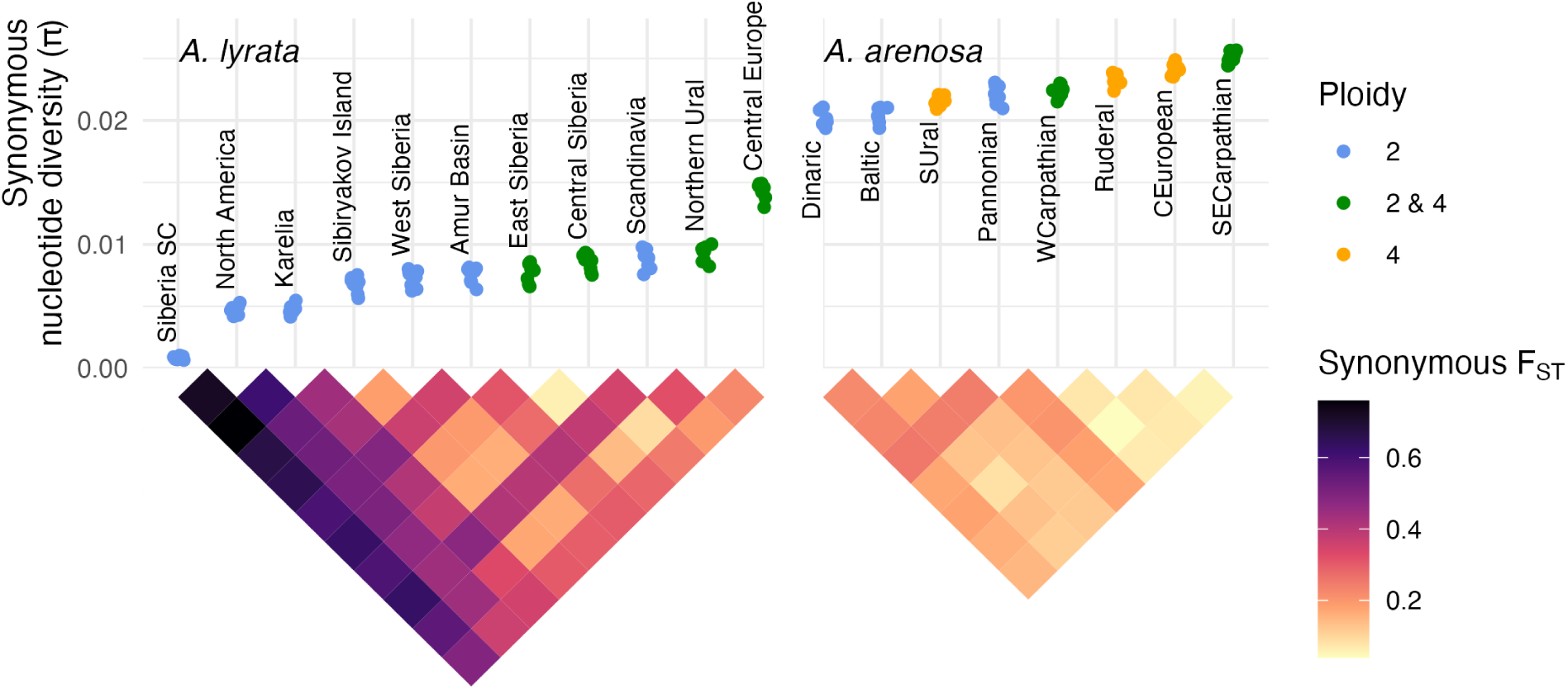
Genome-wide synonymous diversity and differentiation in the *A. lyrata* (a) and *A. arenosa* (b) datasets, calculated on four-fold degenerate sites. Multiple points for each lineage show π□ values for different chromosomes. Heatmaps below show pairwise lineage differentiation based on F_ST_ between the lineages in *A. lyrata* (a) and *A. arenosa* (b). “Siberia SC” marks the self-compatible Siberian *A. lyrata* lineage.

Within *A. lyrata*, synonymous diversity varies dramatically among lineages, supporting a source–colonization model with Central Europe as the primary reservoir of variation (Clauss and Mitchell-Olds 2006; Ross-Ibarra et al. 2008; Ansell et al. 2010; Schmickl et al. 2010; Pyhajarvi et al. 2012). For diversity and divergence analyses, we grouped 895 *A. lyrata* samples into lineages based on genetic structure, excluding 15 individuals highly admixed with *A. arenosa* (fig. 1a, Supplementary Data 1). Central European *A. lyrata* is more diverse (mean π□ ≈ 1.4%) than any other *A. lyrata* lineage, although still less diverse than all *A. arenosa* lineages. In contrast, self-compatible Siberian *A. lyrata* represents a distinct lineage with extremely low diversity (π□ ≈ 0.1%). Four of the five most diverse *A. lyrata* lineages (Central Europe, Northern Ural, West Siberia, Central Siberia) contain all known tetraploid populations. Consistent with its inferred evolutionary history, *A. lyrata* shows strong population structure, with lower within-lineage diversity and higher between-lineage differentiation (F_ST_) than *A. arenosa* (fig. 4). The highest differentiation is found between the Siberian self compatible lineage (“Siberia SC” in the figure) and the rest of the species complex, reaching the F_ST_ of 75.8%.

In contrast to *A. lyrata, A. arenosa* exhibits uniformly high genetic diversity across its range, with the highest estimates in tetraploid lineages (Central European and Ruderal) and mixed ploidy lineages (Western and Southeastern Carpathian; fig. 1, fig. 4a). Leveraging dense population-level sampling, we calculated synonymous diversity and divergence statistics in 76 populations comprising at least five individuals (630 individuals in total, results are in Supplementary Data 2). Population-level Tajima’s *D* ranged from-0.436 to 0.922 (mean = 0.246), indicating no strong deviations from neutrality. Mean within-population synonymous diversity (π□) was 2.0%, varying significantly among lineages and between ploidies (1.8% / 2.1% on average for diploid / tetraploid populations, respectively; two-sided t-test, p-value < 10^-9^). Lineage differentiation (F_ST_) is not as variable as in *A. lyrata*, with the highest F_ST_ (25.2%) between exclusively diploid lineages Pannonian and Dinaric (fig. 4c).

### Patterns of interspecific gene flow in *A. lyrata* populations

We also investigated relationships among lineages based on the patterns of allele sharing using D-statistics (Patterson et al. 2012). We identified excessive allele sharing in the western *A. lyrata* populations (Central Europe, British Isles and Scandinavia) with *A. arenosa*, *A. halleri* and *A. croatica* relative to Siberian and North American *A. lyrata* lineages (fig. 5) (CR, AA/AH/AC;AL_pop1, AL_pop2, where AL = *A. lyrata*, ALP = *A. lyrata* ssp. *petraea*, ALL = *A. lyrata* ssp. *lyrata*, AA = *A. arenosa,* AH = *A. halleri*, AC = *A. croatica*, CR = *C. rubella*). As *A. arenosa* and *A. croatica* are closely related, these results may stem from shared rather than independent gene flow events. Since the interspecific excess of allele sharing was not observed when Siberia and North America *A. lyrata* lineages were compared (fig. 5), gene flow likely occurred from *A. arenosa* to *A. lyrata* ssp. *petrea* after the western and eastern lineage split assuming closer phylogenetic relationships within sub-species. We also observed excess allele sharing with *A. arenosa* in Central Europe relative to the British Isles populations (fig. 5) which indicates within-Europe differences in the strength of *A. arenosa*-*A. lyrata* interspecific gene flow. These results demonstrates additional complexity in the population history of *Arabidopsis* species where gene flow between species has been observed (Novikova et al. 2016) particularly between the polyploid lineages (Jorgensen et al. 2011; Marburger et al. 2019; Bohutínská et al. 2024) that has also contributed to local adaptation (Arnold et al. 2016). Although a strong hybridization barrier has been previously reported between the diploid *A. arenosa* and *A. lyrata* (Lafon-Placette et al. 2017) our results suggest relatively recent interspecific gene flow.

**Fig. 5.**
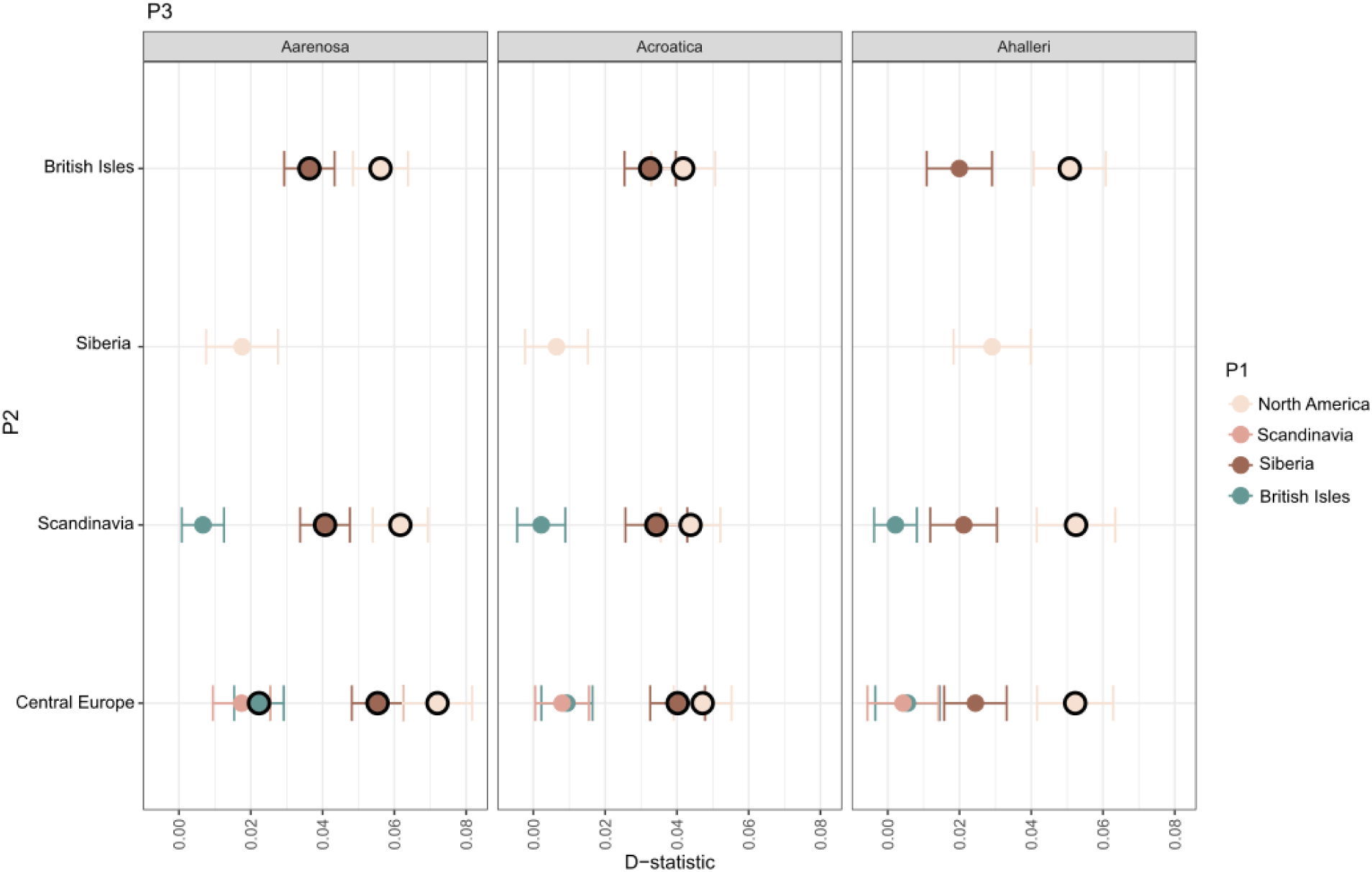
Between species allele sharing in *A. lyrata* populations. D-statistics testing for interspecific allele sharing between different *A. lyrata* populations and *A. arenosa*, *A. halleri* and *A. croatica*. The significant test statistics (Z-value > 3) are highlighted in bold. The error bars show block jackknife standard errors. Positive D indicates increased allele sharing between P2 and P3.

### Genome-wide analysis reveals latitude-associated variants in *A. lyrata*

To test the power of the dataset for identifying environmental associations with genotypes, we selected the Eastern Siberian lineage of *A. lyrata*, which spans the Lena River basin fiig. 2,3) due to its dramatic latitudinal gradient. The variant selection based on sets of individuals (or loci) can be done using the Variant Selector tool at arabidopsyslyrata.org. As an environmental variable, we chose latitude, which correlates with day length and, in our case, temperature (Pearson-correlation-0.93) We conducted a population structure-aware genome-wide association study (GWAS) on 124 individuals belonging to the Eastern Siberian lineage of *A. lyrata* (fig. 1a, fig. 3a). Using a 1%-permutation-based significance threshold (p< 1.61e-12), we identified 44 associated SNPs distributed across the genome (fig. 6a; Supplementary Data 3). Of these, 31 SNPs were located within annotated gene models of the *A. lyrata* NT1 reference genome, corresponding to 20 unique genes (fig. 6a; supplementary fig. S7). Seven of these genes have been functionally characterized in *A. thaliana*. These include **FT**, a photoperiod-sensitive integrator of flowering time (Turck et al. 2008); **RWA4 (AT1G29890)** and **XAPT1 (AT1G68470)**, which process the secondary cell wall polysaccharide xylan and effect the firmness of the cell wall (Lee et al. 2011); and **CARB (AT1G29900; VEN3)**, which, together with VEN6, encodes subunits of carbamoyl phosphate synthetase required for arginine biosynthesis, with mutants showing green veins and pale lamina (Mollá-Morales et al. 2011). We also identified **ACO4 (AT1G13280)**, involved in jasmonic acid biosynthesis (Stenzel et al. 2003) and thus in abiotic and biotic stress signaling (Ghorbel et al. 2021), and the multifunctional **ASA1 (AT3G02260; also BIG/TIR3/LPR1/UMB1/CRM1/DOC1)**, a regulator of auxin transport (Gil et al. 2001) whose mutants exhibit altered root (Ruegger et al. 1997; López-Bucio et al. 2005) and inflorescence morphology (Yamaguchi et al. 2007), delayed flowering (Kanyuka et al. 2003) possibly due to upregulation of the vernalization-sensitive flowering repressor *FLC (Ebine et al. 2012)*. Finally, **ATIPS1 (AT4G39800; MIPS1)** encodes a widely expressed l-*myo*-inositol-1-phosphate synthase (Donahue et al. 2010) that regulates myo-inositol production (Latrasse et al. 2013), contributing to processes such as programmed cell death (Meng et al. 2009) and photoperiod measurement (de Cássia Monteiro Batista et al. 2024).

**Fig. 6.**
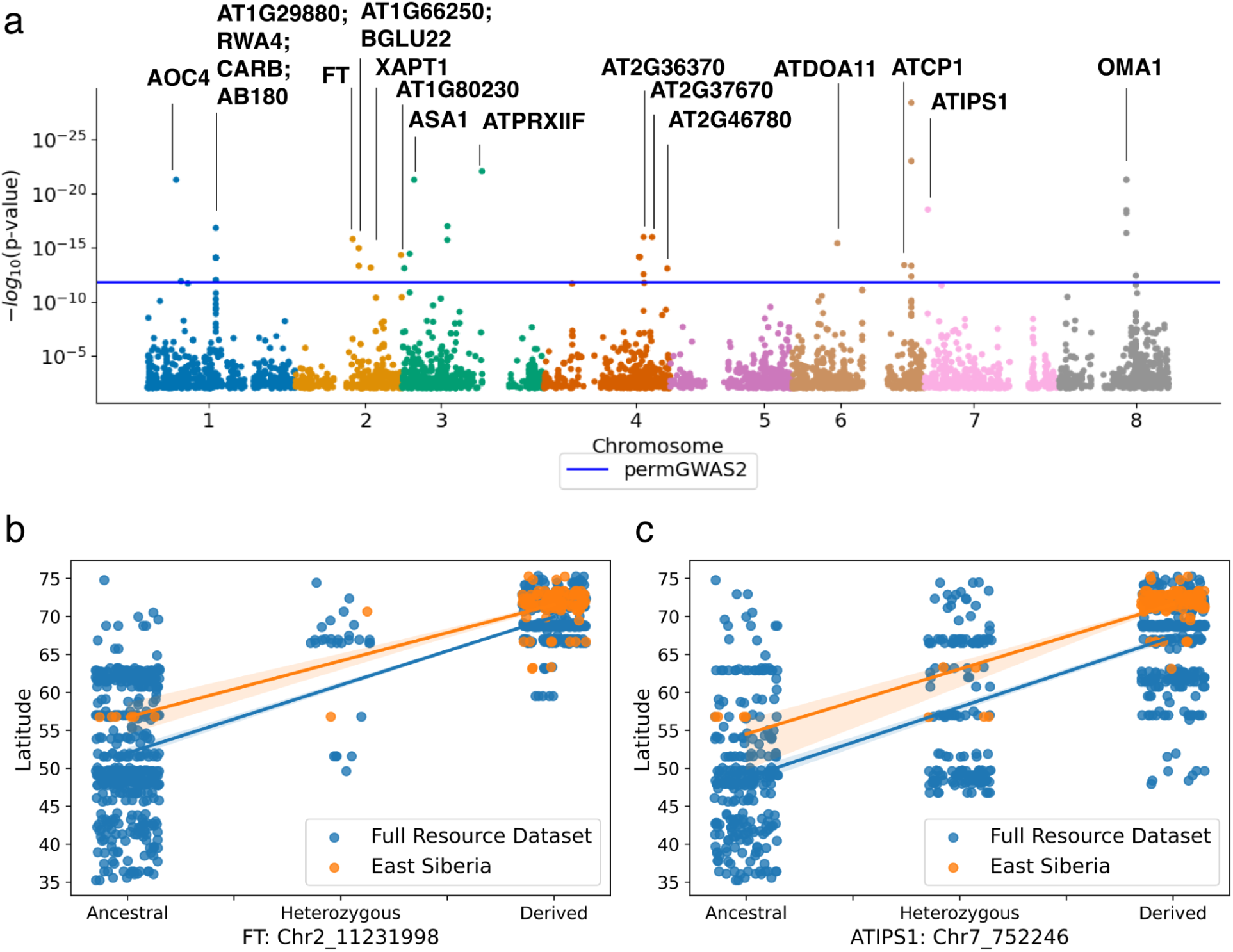
(a) GWAS on latitude on 124 individuals of Eastern Siberian *A. lyrata*. Manhattan plot of genotype associations with latitude; horizontal line represents a 1%-permutation-based significance threshold. Significant peaks are annotated by the underlying gene model. Linear regression of latitude and *FT* (b) and *ATIPS1* genotypes (c) within Eastern Siberian lineage (orange) and across the entire *A. lyrata* dataset (blue). The derived allele is associated with higher latitudes compared to the ancestral allele.

Correct timing of reproductive events - such as flowering in plants - is a key for evolutionary success. Photoperiod is one of the focal external factors controlling growth and reproduction in many plant species (Garner and Allard 1920; Lumsden 2002) and it is highly variable across latitudes. Hence, to time flowering properly using day length cues, plants must set their critical daylength in colonization of high latitudes. From this perspective, among the associated gene variants identified in this study, *FT* and *ATIPS1* are of particular interest due to their central role in photoperiodic control of development (Gendron and Staiger 2023). *FT* is a key flowering time input integrator that collects information from multiple external and internal sensing pathways in *A. thaliana* (Kim 2020). After expression induction in leaves, the protein product of *FT* is transported to meristem where it induces flowering (Corbesier et al. 2007). *FT* homologs have conserved role in day length mediated growth regulation – such as in growth-dormancy fluctuations - in many plant species across the flowering plant phylogeny (Böhlenius et al. 2006; Hayama et al. 2007; Li et al. 2024). Recent investigation in *A. thaliana* revealed that under long-day conditions, reproductive and vegetative growth are controlled independently, with *ATIPS1* acting as a central regulator of the vegetative growth (Wang et al. 2024). Comparing the state of the significantly associated alleles to the outgroups by checking the individual SNPs and the entire CDS (supplementary fig. S7, S8), we found that the Northern haplotype for both *FT* and *ATIPS1* is derived and *A. lyrata* specific.

Recognition of the central role of *FT* in flowering time control, particularly in the photoperiodic pathway, has already early granted its recognition as a strong candidate gene for conferring latitudinal adaptation (Turck et al. 2008). Indeed, genetic variation in the orthologs of *FT* have repeatedly been associated to confer latitudinal adaptation in multiple systems. For example, variation in flowering time is associated with different naturally occurring haplotypes of *FT* ortholog in soybean (*Glycine max*), *GmFT2b* (Chen et al. 2020). An early flowering haplotype, Hap3, was more common in higher latitudes while two other haplotypes Hap1 and Hap4 could not flower in high latitude environments suggesting that the variation is *FT2b* has allowed high latitude adaptation in this species (Chen et al. 2020). Furthermore, variants in *FT* ortholog were also identified potentially contributing to latitudinal adaptation in rice bean (*Vigna umbellata*) (Guan et al. 2022).

Our findings of latitudinal differentiation in *FT* and *ATIPS1* alleles could be due to orchestrated change in the regulation of photoperiodic growth and reproduction in the high-latitude *A. lyrata* populations. Previous investigations of the Northwest Eurasian *A. lyrata* found no evidence for selective sweep or local adaptation in *FT,* while such signatures were observed in other photoperiodic pathway flowering time genes upstream of *FT,* such as pleiotropic photoperiodic pathway gene *GIGANTEA (GI)* (Mattila et al. 2016) and light quality sensing *PHYTOCHROME A* (*PHYA*) (Toivainen et al. 2014; Mattila et al. 2016). Together, this suggests independent adaptation to higher-latitude in North and East Eurasia, targeting different core components of photoperiodic regulation.

### Outlook

By providing convenient open access to genome-wide variation in *A. lyrata* and *A. arenosa*, we advance population genomics toward the FAIR principles (Wilkinson et al. 2016): Findable, Accessible, Interoperable, and Reusable. The access to pre-processed variant data also significantly reduces human-and computational-time demands as it allows avoiding the most demanding and often redundant steps involving gathering primary data and metadata from nucleotide repositories, raw read pre-processing, mapping and variant calling. The scale and resolution of this dataset allow investigation of how genomic architecture, including recombination landscapes, structural variation, and ploidy, shapes adaptation to environmental gradients, as well as dissection of the origins of environmentally associated alleles through standing variation, introgression, and parallel evolution. The possibility to seamlessly integrate, for example gene expression and phenotypic information with this world-wide diversity data has a potential to support a variety of investigations ranging from gene level to the entire genome-wide approaches across both species.

## Acknowledgements

We thank Jose Reza and Ümit Seren for providing IT support for the website development. We acknowledge the European Union ERC HOW2DOUBLE, 101041354 to P.Y.N., ERC DOUBLE ADAPT 850852 to F.K., and ERC HOTSPOT 679056 to L.Y., the Research Council of Finland [decision number 360310] to T.M.M, the joint Deutsche Forschungs gemeinschaft (DFG) and GACR program—project number 490698526 to P.Y.N. and 22-29078K to F.K. and the exchange program by DAAD fund 57601834 for promoting scientific exchange between the groups of P.Y.N. and F.K. M.B. was supported by the European Union’s research and innovation programme under the Marie Skłodowska-Curie grant agreement 101062703-CONstrainCONverge. The results of this project LL2317 were obtained with the financial contribution of the Ministry of Education, Youth and Sports as part of the targeted support of the ERC CZ program. T.M.M. and J.F. acknowledge the e-INFRA CZ project (ID:90254), supported by the Ministry of Education, Youth and Sports of the Czech Republic and CSC – IT Center for Science, Finland, for providing computational resources. We also thank colleagues who shared with us plant material for sequencing: Alexey Seregin, Alexey Penin, Heidi Serra and Team Evo-Eco. The views and opinions expressed are however those of the author(s) only and do not necessarily reflect those of the European Union, Research Council of Finland or the European Research Council Executive Agency. Neither the European Union nor the granting authority can be held responsible for them.

## Methods

### Plant material

To minimise taxonomic bias in our inference, we consider each major lineage as a single species, i.e. *A. arenosa* s.l. and *A. lyrata* s.l. (*sensu lato*, i.e. in a broad sense). That means we included all taxonomic entities traditionally associated with each of these broadly recognised species, including those that were previously shown outside of major genetic grouping within each species. Specifically, the following morphologically-delimited taxa are included within *Arabidopsis arenosa* (L.) Lawalrée as previous genetic studies had not supported this separation (e.g. (Kolář, Fuxová, et al. 2016; Knotek et al. 2020; Padilla-García et al. 2023)): *A. arenosa* subsp. *borbasii* (Zapałowicz) O’Kane & Al-Shehbaz, *A. carpatica* nom. prov., *A. nitida* nom. prov., *A. neglecta* (Schultes) O’Kane & Al-Shehbaz, *A. petrogena* (A. Kern) V.I. Dorof. Instead, we recommend referring to the genetic lineages (and possibly also ploidy cytotypes if such data are available) presented here. In contrast, *Arabidopsis croatica* (Schott) O’Kane & Al-Shehbaz is kept as separate species due to its distinct phylogenetic position shown by both genome-wide and plastome markers (Kolář, Fuxová, et al. 2016; Monnahan et al. 2019).

In *A. lyrata*, classification based on morphological data also disagrees with genetics, the latter often revealing even misclassified allotetraploid *A. kamchatica* as diploid *A. lyrata* in Eastern Eurasia and in North America. Moreover, there are several subspecies classifications with different perceptions of the same subspecies name, as well as different subsets of subspecies. (Al-Shehbaz and O’Kane 2002) distinguished three subspecies: *A. lyrata* subsp. *lyrata* in North America and NE European Russia, *A. lyrata* subsp. *petraea* (L.) O’Kane & Al-Shehbaz in Europe but also in Siberia and Far East, and *A. lyrata* subsp. *kamchatica* in North America, Eastern Siberia and Far East. In the checklist of Panarctic flora, Elven (ed.) 2007 (cited by (Schmickl et al. 2010)) did not include *A. lyrata* subsp. *lyrata* in *A. lyrata* and considered it all *A. arenicola* (Richardson ex Hook.) (Al-Shehbaz and O’Kane 2002) They distinguished three subspecies: *A. lyrata* subsp. *petraea* in Europe, *A. lyrata* subsp. *septentrionalis* (N.Busch) D.F.Murray & Elven in Siberia and *A. lyrata* subsp. *umbrosa* (Turcz. ex Steud.) D.F.Murray & Elven in Far East, Alaska and Canada. Here we collected subspecies identification as indicated in the source publications or in herbarium labels for plants from Moscow University herbarium. On the PCA (supplementary fig. S5), we can see that *A. lyrata* subsp. *lyrata* forms a cluster together with *A. arenicola*, and this cluster encompassing all North American accessions is clearly separated from the rest of the species. Samples from Central Europe and Scandinavia also form a separate cluster and are always identified as *A. lyrata* subsp. *petraea*. However, samples identified as *A. lyrata* subsp. *petraea* in Russia overlap with other identifications as *A. lyrata* subsp. *septentrionalis* and *A. lyrata* subsp. *umbrosa* without a clear separation between them. We thus suggest treating all samples as *A. lyrata* complex without separation of subspecies or microspecies (*A. arenicola*) and rather refer to the genetic lineages presented here.

Locality details on both previously and newly sequenced accessions are reported in Supplementary Data 1 and 2.

### DNA sequencing

For the short-read sequencing libraries preparation we used about 1 cm^2^ of leaf tissue. For *A. lyrata* samples, genomic DNA was isolated with the “NucleoMag© Plant” kit from Macherey and Nagel (Düren, Germany) on the KingFisher 96Plex device (Thermo) with programs provided by Macherey and Nagel. Libraries were pooled and sequenced on the NovaSeq 6000 S4 flow cell Illumina system. For *A. arenosa* samples, genomic DNA was extracted from silica-gel-dried leaves using the sorbitol extraction method (Štorchová et al. 2000) and then purified using AMPure XP (Beckman Coulter Inc., Brea, California, USA). Protocol LITE of Perez-Sepulveda et al. 2021 was used for the library construction (Perez-Sepulveda et al. 2021). The libraries were sequenced in 300 cycles (2 × 150 bp paired-end/PE reads) on the Illumina NovaSeq platform.

### Read mapping and variant calling

*A. lyrata* reads obtained both from sequencing and from available databases were processed as described in (Kolesnikova et al. 2023). Reads were trimmed using BBDuk (Bushnell 2014), mapped on NT1 *A.lyrata* genome (Kolesnikova et al. 2023) using bwa mem (Li and Durbin 2009), processed to remove the duplicated reads with samtools (Danecek et al. 2021), and genotypes were called with GATK (McKenna et al. 2010). GATK parameter--includeNonVariantSites was used to keep all sites in the vcf file. The vcf file was merged with a file used in (Scott et al. 2025) using bcftools merge (Danecek et al. 2021). This gave us a raw VCF file with both variant and invariant unfiltered sites (210,861,218 total records, out of which 85,109,507 were SNPs and 38,042,130 short insertions/deletions) which is accessible through the resource website (file: All_lyrata_final_allpos.vcf.gz). For vcf of biallelic SNPs GATK hard filtering was used in addition to filtering only biallelic SNPs and filtering by minimum coverage of 3 (available as file: All_lyrata_final_allpos_bial.vcf.gz). This vcf was used to filter 4-fold degenerate sites (available as file: All_lyrata_final_allpos_4fds.vcf.gz). Site annotation was performed using Degenotate (Mirchandani et al. 2024) on Helixer (Stiehler et al. 2021) annotation. Only gene models with CDS longer than 300 nucleotides were kept, filtering was performed with AGAT (Stiehler et al. 2021; Dainat et al. 2024).

We gathered short-read sequencing data of *A. arenosa* from all available published datasets (Hollister et al. 2012; Yant et al. 2013; Arnold et al. 2016; Novikova et al. 2016; Monnahan et al. 2019; Preite et al. 2019; Bohutínská et al. 2021; Konečná et al. 2021; Weitz et al. 2021; Bohutínská et al. 2024) and complemented them with additional sequencing. In total, we gathered sequences for 736 accessions: 723 samples of *A. arenosa* from 150 natural populations (N individuals range = 1-16, mean = 4.9, median = 5, 76 populations with >= 5 individuals) and 13 additional samples of its sister species *A. croatica* available as an outgroup for analyses (Summarized in Supplementary Data 2). Population details are summarized in the Supplementary Data 6.

We used the snakemake pipeline designated to handle mixed ploidy datasets following our analysis best practices (Bohutínská et al. 2024) to process the raw data (https://github.com/jgerchen/polyploid_variant_calling). Specifically, we first used trimmomatic-0.38 to remove adaptor sequences and low-quality base pairs (<15 PHRED quality score). Trimmed reads > 50 bp were mapped to reference genome assembly of diploid *Arabidopsis arenosa* from the Western Carpathian lineage (corresponding to our population PUP; Bramsiepe et al. 2023) by bwa--0.7.17 (Li and Durbin 2009) with default settings. Duplicated reads were identified by picard-2.8.1 (https://github.com/broadinstitute/picard) and discarded together with reads that showed low mapping quality (<20). Afterwards we used GATK v.4.0 to call and filter reliable variants and invariant sites according to recommended best practices. Namely, we used the HaplotypeCaller module to call variants per individual using the ploidy = 2 and 4 set for each individual according to the prior ploidy inference information (see Supplementary Data 2 for details). Then, we jointly called variants and invariants across all individuals by module GenotypeGVCFs. This gave us a raw VCF file with both variant and invariant unfiltered sites (148,859,087 total records, out of which 78,843,205 were SNPs and 31,464,040 short insertions/deletions) which is accessible through the resource website (file: arenosa_full.vcf.gz).

We retained biallelic SNPs and further filtered these sites following GATK best-practice recommendations: Quality by Depth (QD) < 2.0, FisherStrand (FS) > 60.0, StrandOddsRatio (SOR) > 3.0, RMSMappingQuality (MQ) < 40.0, MappingQualityRankSumTest (MQRS) < −12.5, and ReadPosRankSum < −8.0. Genotypes with read depth (RD) < 4 per individual were set to no-call (∼ missing data). The resulting dataset comprised 736 individuals and 16,933,635 biallelic SNPs (file: arenosa_biallelic.vcf.gz). Finally, we selected only 4-fold degenerate sites (both variants and invariants) from the raw VCF file and applied the same filtration criteria as described above for the biallelic SNPs (736 individuals, 4,304,682 records out of which 2,849,754 were SNPs). This file is also available at the resource website (file: arenosa_4folds.vcf.gz). For analyses of population structure and diversity, we used a subset of 3,451,486 4-fold degenerate sites (2,002,056 biallelic SNPs and 1,449,430 filtered invariant positions) from this dataset.

### Gene orthology search

Correspondence of Helixer gene models to known genes of *A. thaliana* and *A. lyrata* was found as a major vote of the following methods. First, we overlapped Helixer and liftoff annotations (Shumate and Salzberg 2021; Stiehler et al. 2021; Dainat et al. 2024) using GffCompare (Pertea and Pertea 2020). In addition, we used GENESPACE (Lovell et al. 2022) to find orthologs in MN47 *A. lyrata* (Hu et al. 2011; Lovell et al. 2022) and TAIR10 *A. thaliana* genomes. With this approach, we only considered syntenic genes, and a single correspondence with maximum bitscore was extracted for each gene.

To assign orthology relationships between *A. arenosa* and *A. thaliana* genes, we ran OrthoFinder v.2.5.4 (Emms and Kelly 2015; Emms and Kelly 2019) with proteomes of 22 Brassicaceae species with reference genomes and annotations of suitable quality. Additionally, we ran also BLAST reciprocal best hits analysis (Cock et al. 2015) with default values (minimally 70% identity over the aligned region, which should cover at least 50% of the query sequence) and simple BLAST search (blastp 2.16.0+ (Camacho et al. 2009)) of *A. arenosa* proteome against *A. thaliana* proteome. We assigned *A. thaliana* orthologues (or other kinds of homologues) to the genes of *A. arenosa* in one of five ways, using the next step only if no *A. thaliana* gene was assigned in the previous one: (1) Orthologues (from “Orthologues” directory) produced by OrthoFinder, (2) BLAST reciprocal best hits, (3) genes from N0 phylogenetic hierarchical orthogroups produced by OrthoFinder, (4) genes from broader orthogroups (from “Orthogroups” directory) produced by OrthoFinder and (5) best hits (based on bitscore) from simple BLAST search (E-value minimally 0.1, identity over the aligned region minimally 70%, aligned region should cover at least 50% of the query sequence). The scripts and details describing orthology assignments for *A. arenosa* are available at https://github.com/mduchoslav/Brassicaceae_orthology.

Resulting orthology tables are available as Supplementary Data 4 (*A. lyrata*) and 5 (*A. arenosa*). The genome browser is also searchable by TAIR10 names.

### *Arabidopsis* genus gene tree - species tree reconciliation analysis

The raw reads of diploid accessions from a representative set of lineages across the entire genus served as input for ParalogWizard (Ufimov et al. 2022). ParalogWizard requires specific references (one with separated exons and another with concatenated exons). To build these references, we extracted each annotated mRNA from the *A. lyrata* reference genome (Hu et al. 2011) both as individual exon fragments and as exon-concatenated sequences using 2ex (https://github.com/rufimov/2ex/tree/main) based on the annotation with RNA-Seq-assisted refinement (Rawat et al. 2015) (retrieved from Phytozome (Goodstein et al. 2012)).

All *A. lyrata* mRNAs were then searched against themselves with BLAST v2.16.0+ in blastn mode, retaining high-scoring segment pairs that covered at least 50 % of the query length. Loci for which each transcript produced exactly one hit (i.e. putative single-copy genes) were deemed orthology-ready and passed to ParalogWizard. In ParalogWizard we assembled the reads for each locus and produced multiple-sequence alignments without invoking paralogue detection, so the program generated orthologous contig alignments using MAFFT v7.453 under default settings.

Downstream analysis followed the HybPhyloMaker (Fér and Schmickl 2018) workflow. Two outgroup taxa, *Capsella rubella* (assembly Caprub1_0 GCF_000375325.1) (Slotte et al. 2013) and *Arabidopsis thaliana* (assembly TAIR10.1 GCF_000001735.4) (Swarbreck et al. 2008), were incorporated by extracting their exon sequences with 2ex and bypassing the assembly step. These reference exons were aligned jointly with other contigs, ensuring orthology of all loci for phylogenomic inference. Within every alignment we first deleted taxa exhibiting more than 70 % missing data, after which loci represented in fewer than 75 % of the remaining samples were discarded. Maximum-likelihood gene trees were estimated in RAxML v8.2.4 (Stamatakis 2014) under the GTRGAMMA substitution model with 1 000 rapid bootstrap replicates. The gene trees were used to infer a species tree with ASTRAL-IV v1.20.4.6 (Zhang et al. 2025). The resulting species tree was then dated using treePL v1.0 (Smith and O’Meara 2012). Optimal cross-validation parameters were estimated automatically using a wrapper script (v0.9; available at https://github.com/tongjial/treepl_wrapper/). In addition to the species tree, only gene trees containing the root were retained and dated. As a secondary calibration, the root corresponding to the split between Capsella rubella and Arabidopsis was constrained using a stem age of the tribe Arabidopsideae of 12.2 million years (My) (95% interval: 11.4–13.1 My;(Hendriks et al. 2023)). The dated species and gene trees were visualized using a DensiTree plot generated with the R package phangorn v2.12.1. As only rooted gene trees can be utilized in DensiTree, 712 rooted gene trees were shown. To further assess gene tree incongruence, quartet sampling was performed using quartetsampling v1.3.1 following (Pease et al. 2018).

### Network analysis, population admixture

Between-individual distance matrices were inferred from variant calls at fourfold degenerate sites, multiallelic SNPs and indels excluded, using VCF2Dis v1.55 (Xu et al. 2025). The NeighborNet network was built using SplitsTree v4.19.0 (Huson and Bryant 2006). ADMIXTURE (Gutenkunst et al. 2009) analysis on *A. lyrata* was performed as described in (Gutenkunst et al. 2009; Scott et al. 2024). For *A. arenosa*, we used Entropy (Shastry et al. 2021), a clustering method designed to analyse mixed-ploidy datasets. Specifically, we run the Entropy with default parameters to investigate population genetic structure using LD-pruned 4-fold degenerate sites (10,226 SNPs), with the number of clusters (K) being set from 2 to 15.

### Estimation of nucleotide diversity and divergence

Within-lineage nucleotide diversity (π) and Tajima’s D and between-lineage fixation index (F_ST_, Hudson’s estimator) values were calculated across four-fold degenerate sites with piawka v0.9.0 developmental (https://github.com/novikovalab/piawka; (Scott et al. 2025)). Multiallelic SNPs and indels were excluded from the analysis using bcftools v1.22 (Danecek et al. 2021). Clusters found in population structure analysis were used to delimit lineages. For *A. lyrata*, two additional lineages (Central Siberia and Amur Basin: Supplementary Data 1) were delimited to account for population structure observed in the network.

### Quantification of allele sharing and shared drift

We calculated f3-outgroup and D-statistics (Patterson et al. 2012) using popstats (Skoglund et al. 2015) using only polymorphic sites in pop1 and 2 as well as pop3 and 4 comparisons (--informative flag). The 4-fold degenerate site vcf file was converted to plink tped format using the vcf2plink.py script supplied in the popstats package and the polyploid lineages were excluded. The popstats.py was converted to python3 using python 2to3.

### RNA sequencing and mapping

Short-read RNA-seq data (ERR14294633, (Kolesnikova et al. 2023)) and (PRJNA459481, (Hämälä et al. 2022)) was mapped to the reference using hisat2 (Zhang et al. 2021). For long-read RNA sequencing, we pooled together leaf and flower tissue. Total RNA was isolated with RNeasy Plant kit, Qiagen including a DNase I treatment during isolation followed by RNA quality assessment by capillary electrophoresis on a NanoChip, Agilent Bioanalyser. Sequencing libraries were then prepared according to the protocol “Procedure & Checklist – Iso-Seq™ Express Template Preparation for Sequel® and Sequel II Systems” of the vendor. Finally, IsoSeq sequencing was performed on a single 8 M SMRT cell on a Sequel IIe device at the Max Planck Genome-centre Cologne. Isoseq data was mapped with minimap2 (options:-ax splice-uf-k14-G2k) (Li 2018). Methylation data (Hämälä et al. 2022) was mapped to the reference using Bismark v0.24.2 (Krueger and Andrews 2011).

### Genome-wide Association Studies (GWAS)

We performed a GWAS using a subset of 124 *A. lyrata* individuals classified within the East Siberia admixture group (see Supplementary Data 1). Variant data were filtered from the original VCF file to include only these individuals. SNPs with missing data were excluded. To optimize the genomic inflation factor (λ) towards the ideal value of 1.0 (λ = 1.05 achieved) a minor allele frequency threshold of 10% was applied. For GWAS, genotype data were encoded into an additive count-matrix, representing the number of minor alleles. Association testing was conducted using permGWAS (John et al. 2022), a permutation-based GWAS framework including a kinship matrix to correct for population structure. To correct for multiple testing, 10,000 phenotype permutations were performed, and SNPs were considered significant at a 1% empirical p-value threshold (p < 1.61e-12). Significant associations were visualized using Manhattan plots. Peaks in the plot were annotated if significant SNPs were located within annotated gene models of the NT1 reference annotation. These gene models were further annotated using the corresponding *A. thaliana* gene name, as provided by the TAIR10 database (https://www.arabidopsis.org).

For each significant SNP, a linear regression model between SNP genotype and latitude was computed using the statsmodels interface in Python, based on the same 124 individuals selected for the GWAS. A second linear model was calculated using the same parameters for all *A. lyrata* individuals from the global dataset. The resulting regression lines were visualized together with the SNP genotypes of individual samples, producing one plot per SNP position.

We polarized SNPs as ancestral or derived using outgroup species included in the dataset. When all outgroup species shared the same allele at a given SNP, that allele was inferred as the ancestral state, and the reference NT1 allele was polarized accordingly (e.g., for the FT gene). When outgroup species did not share a single SNP state, we inferred ancestry by reconstructing a gene tree based on the gene’s DNA sequence from all individuals. Gene trees were rooted using *Capsella rubella* as an outgroup and were constructed with IQ-TREE (Nguyen et al. 2015) using 1,000 ultrafast bootstrap replicates and 1,000 SH-aLRT replicates. The resulting tree was visualized with ggtree (Yu et al. 2017).

## Data availability

The raw long read (Iso-seq) RNA data for *A. lyrata* NT1 accession are available on ENA by accession number ERR15526820. Newly whole-genome sequenced samples in this study are available at ENA as bioproject PRJEB49474. Previously available samples used in this study are listed in the Supplementary Data 1 and 2. VCFs are available for selection based on samples and download at https://www.arabidopsislyrata.org.

