## Supplementary material for "A Global Genomic Resource for Outcrossing *Arabidopsis lyrata* and *Arabidopsis arenosa*": SupplenemtaryMaterial

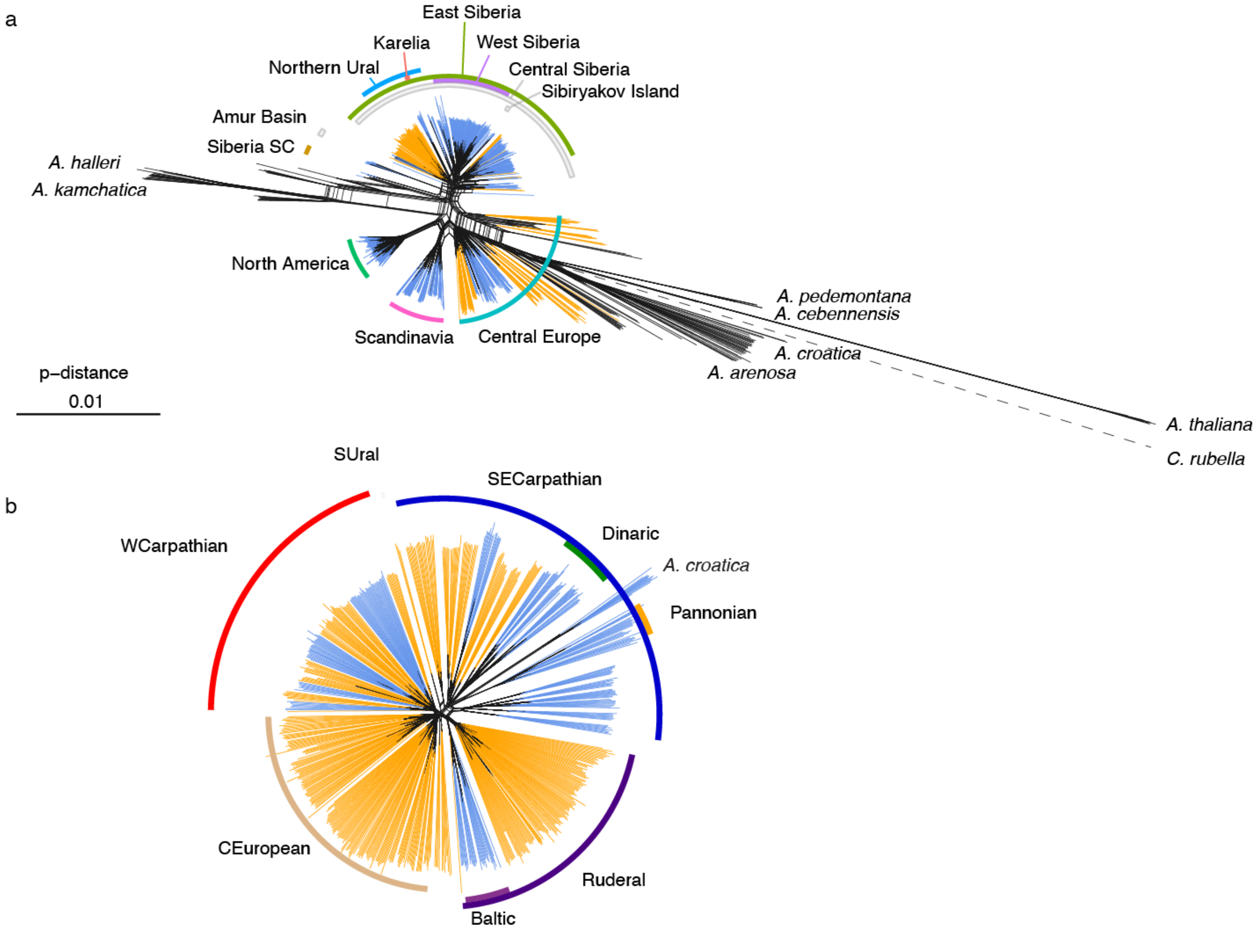


Supplementary fig. 1. Genetic clustering of 1018 samples of *A. lyrata* and other *Arabidopsis* species and outgroup samples included in the same matrix (a) – and clustering of 723 *A. arenosa* and 13 *A. croatica* samples (b) visualised by Neighbor Joining network based on *p*-distances and plotted on the same scale (for fully labelled population network see Supplementary Fig. 4). The analysis is based on synonymous four-fold degenerated SNPs. The blue/orange colour of individual branches corresponds to ploidy (2x/4x), the colour designation of arcs represents the major lineages generally corresponding with the cluster assignments provided in fig. 3. “Siberia SC” marks the self-compatible Siberian *A. lyrata* lineage. Dotted line indicates arbitrary length modified for visualization.

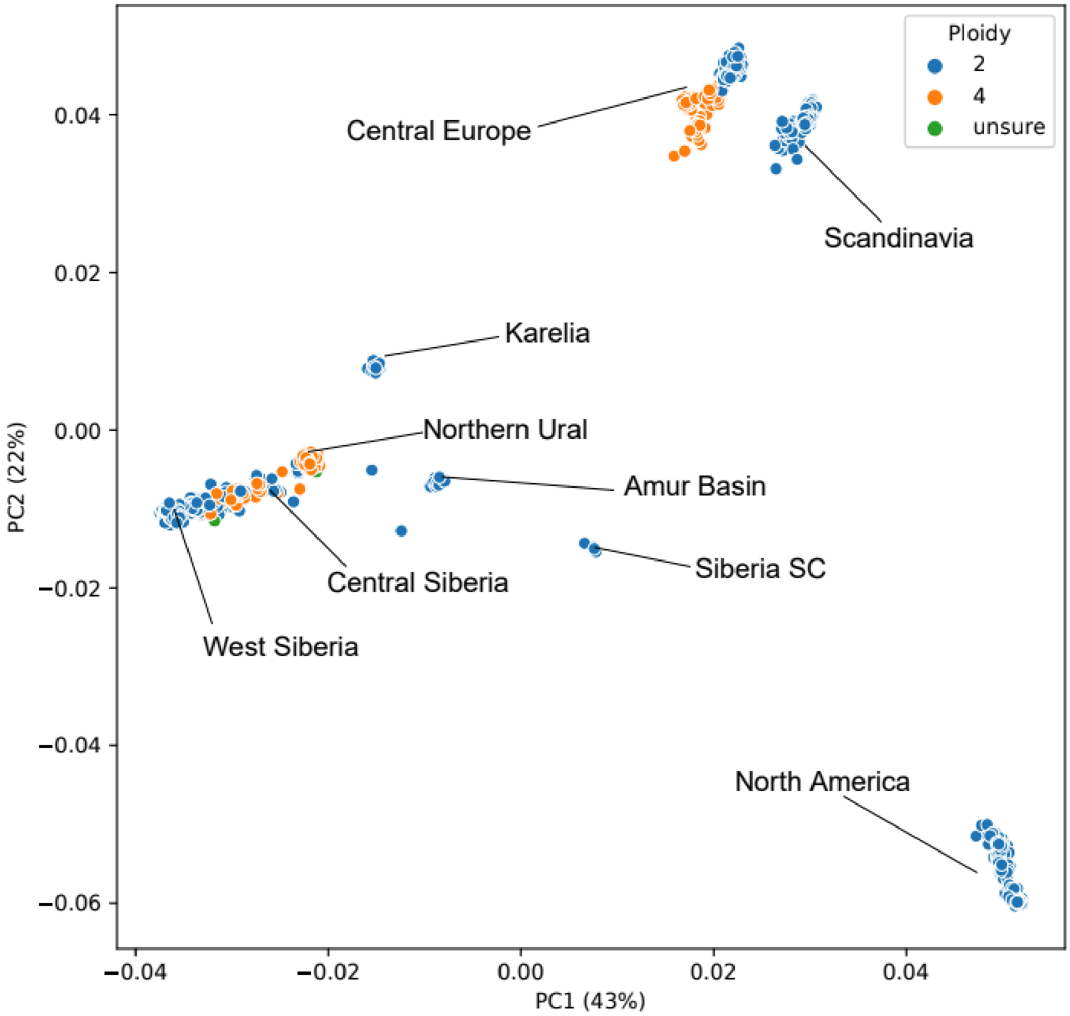


Supplementary fig 2. PCA analysis of the *A. lyrata* samples included in the dataset, the clusters are labeled by the names of the lineages.

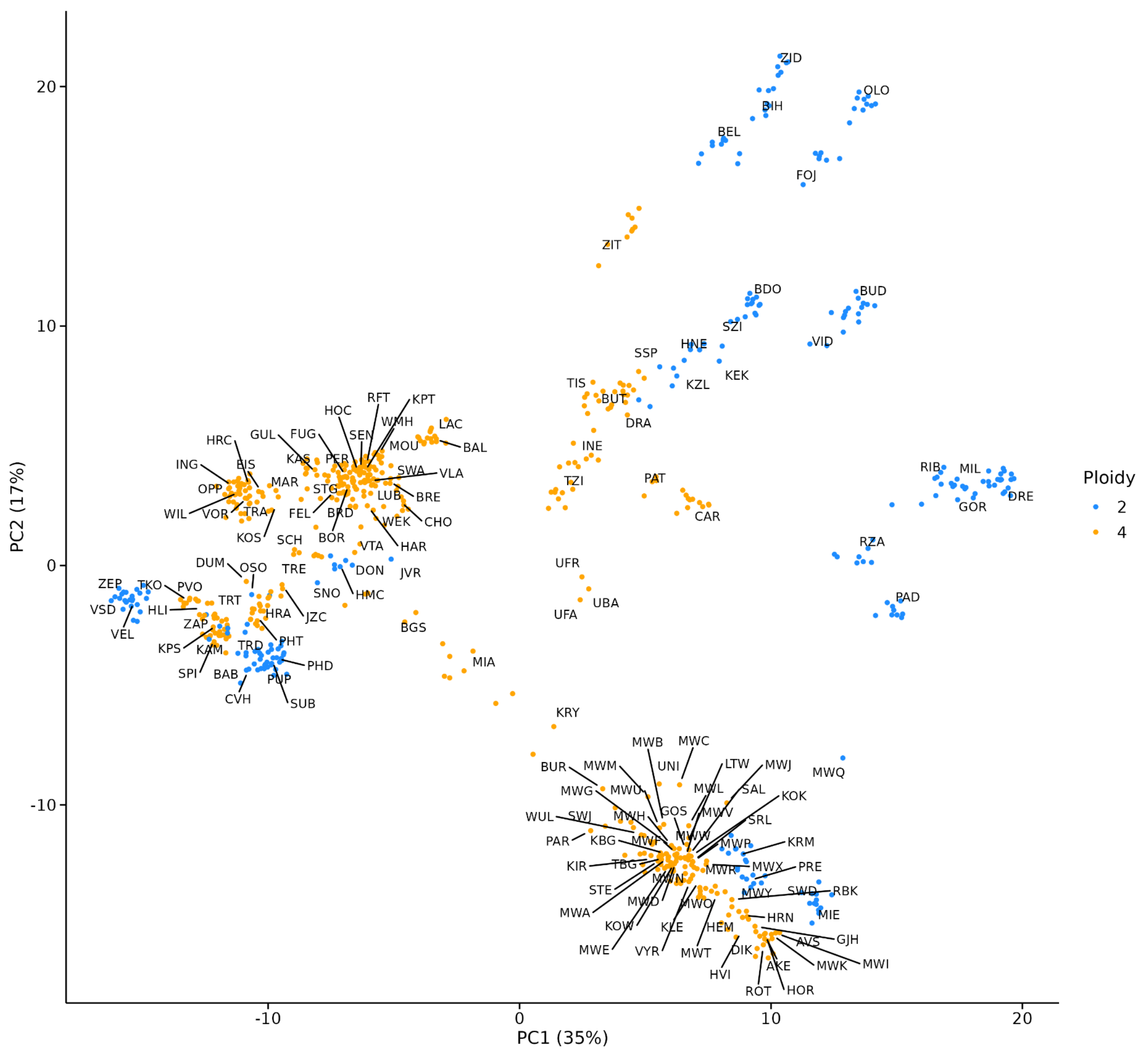


Supplementary fig 3. PCA analysis of the *A. arenosa* samples included in the dataset, the clusters are labeled by the names of the populations.

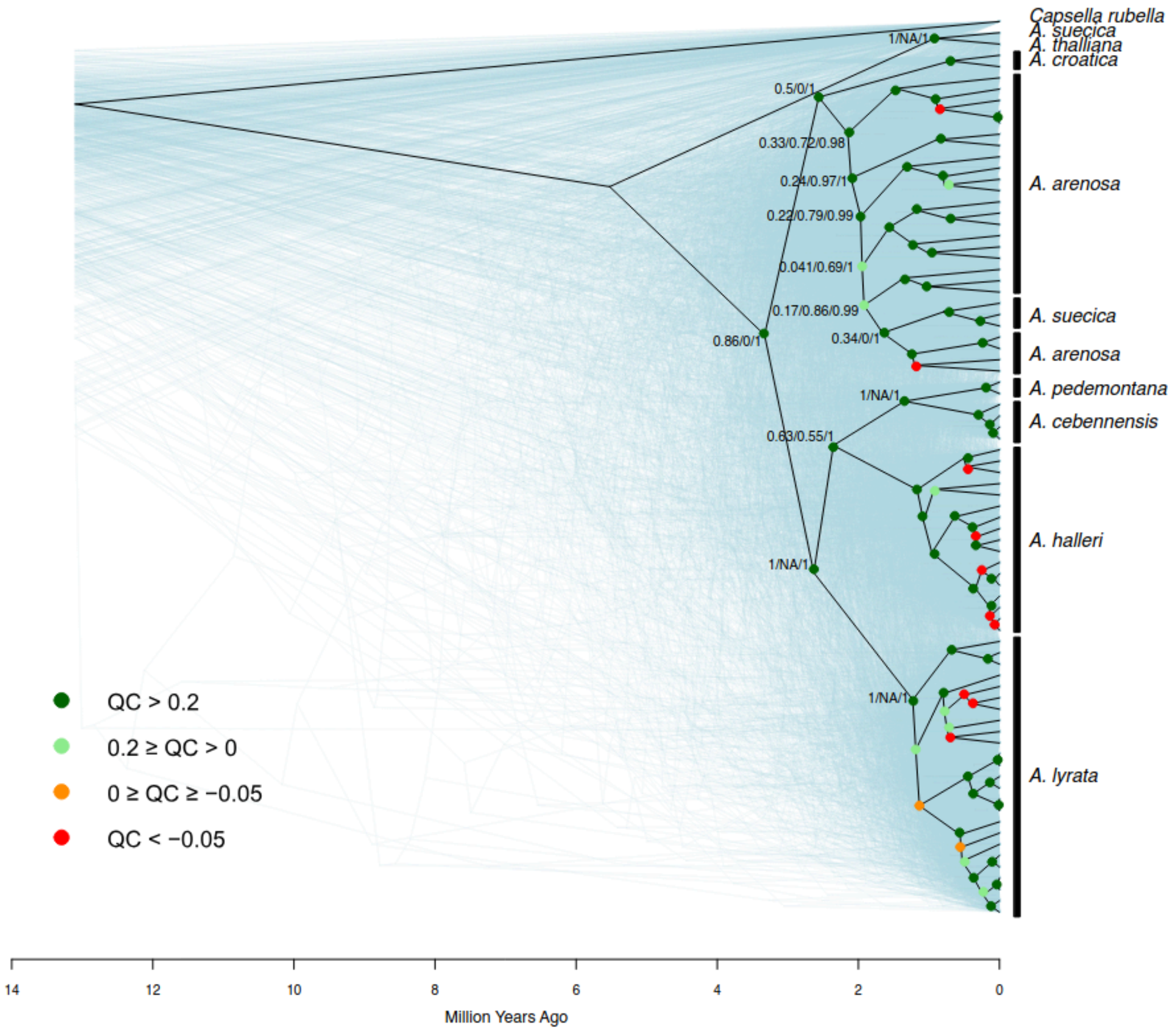


Supplementary fig 4. Consensus phylogenetic tree of *Arabidopsis* genus based on 903 single-copy genes.

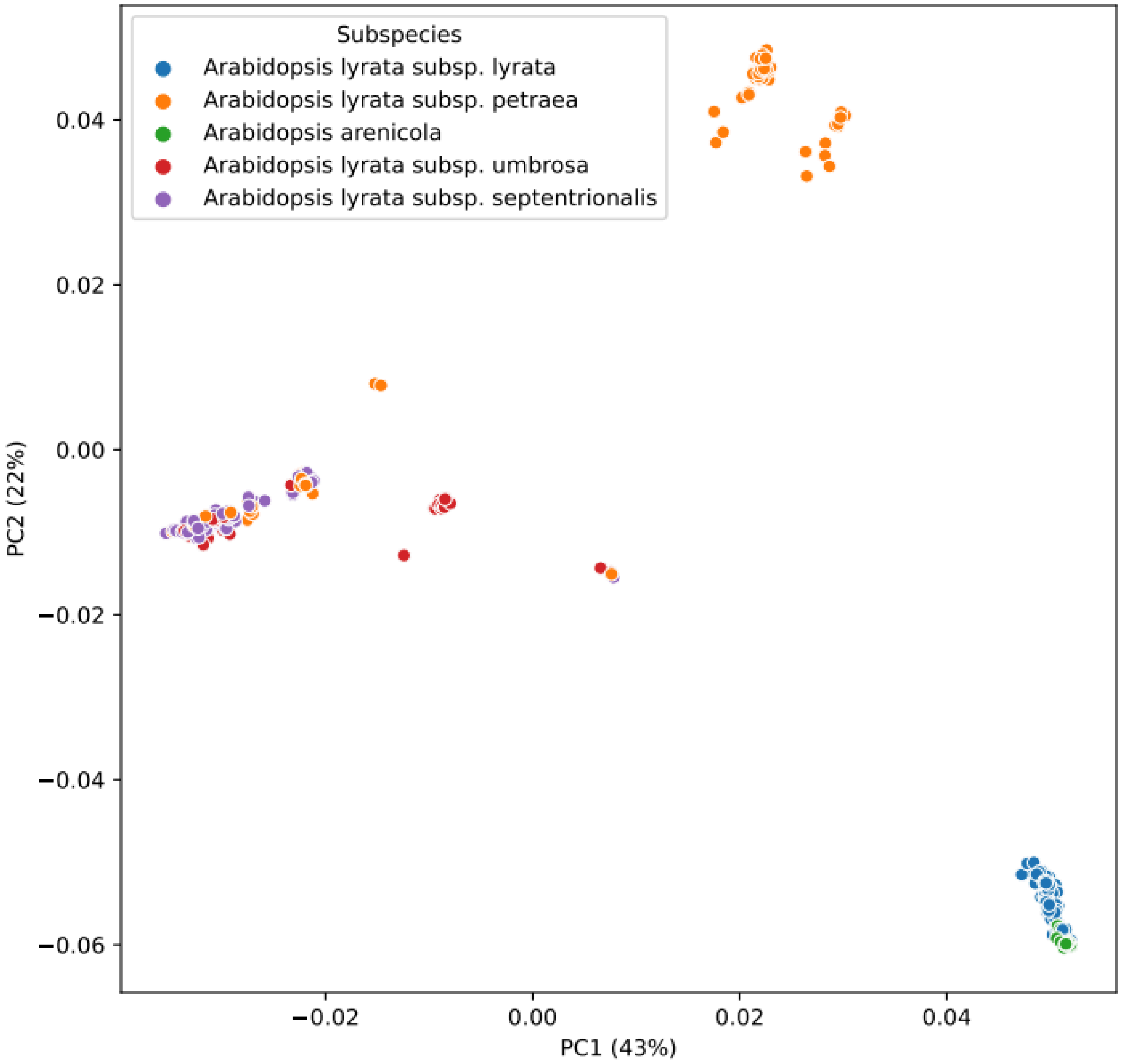


Supplementary fig. 5. PCA analysis of the *A. lyrata* samples included in the dataset, only samples with specified subspecies information are plotted.

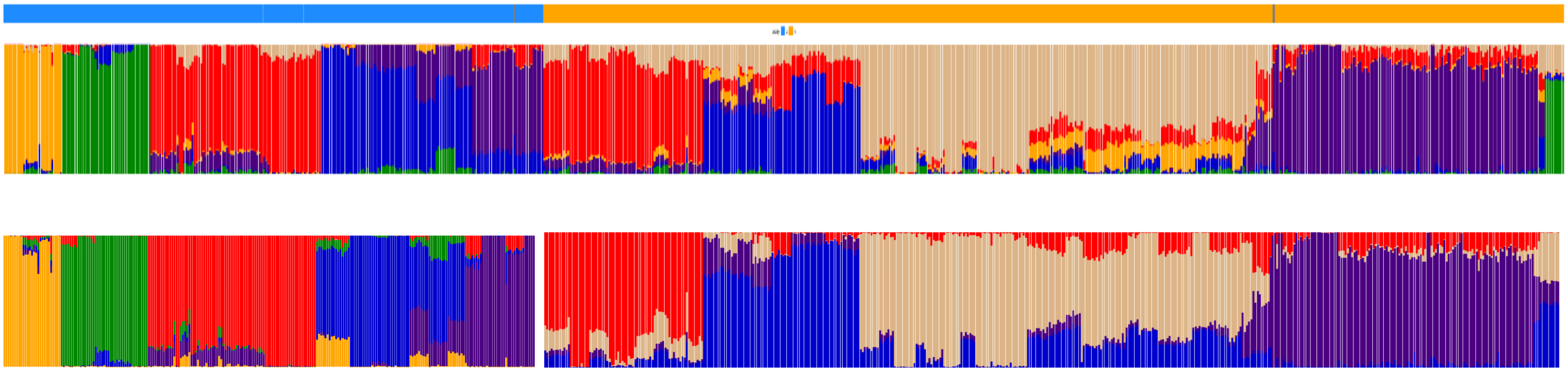

Supplementary fig. 6. Genetic clustering of the 723 *A. arenosa* samples. Individual assignment to groups inferred using Entropy. Upper row: complete dataset (K = 6), lower row, separate analysis of diploids (K = 5) and tetraploids (K = 4). The colour bar on top denotes diploid (blue), tetraploid (orange) and unknown ploidy (gray) individuals.

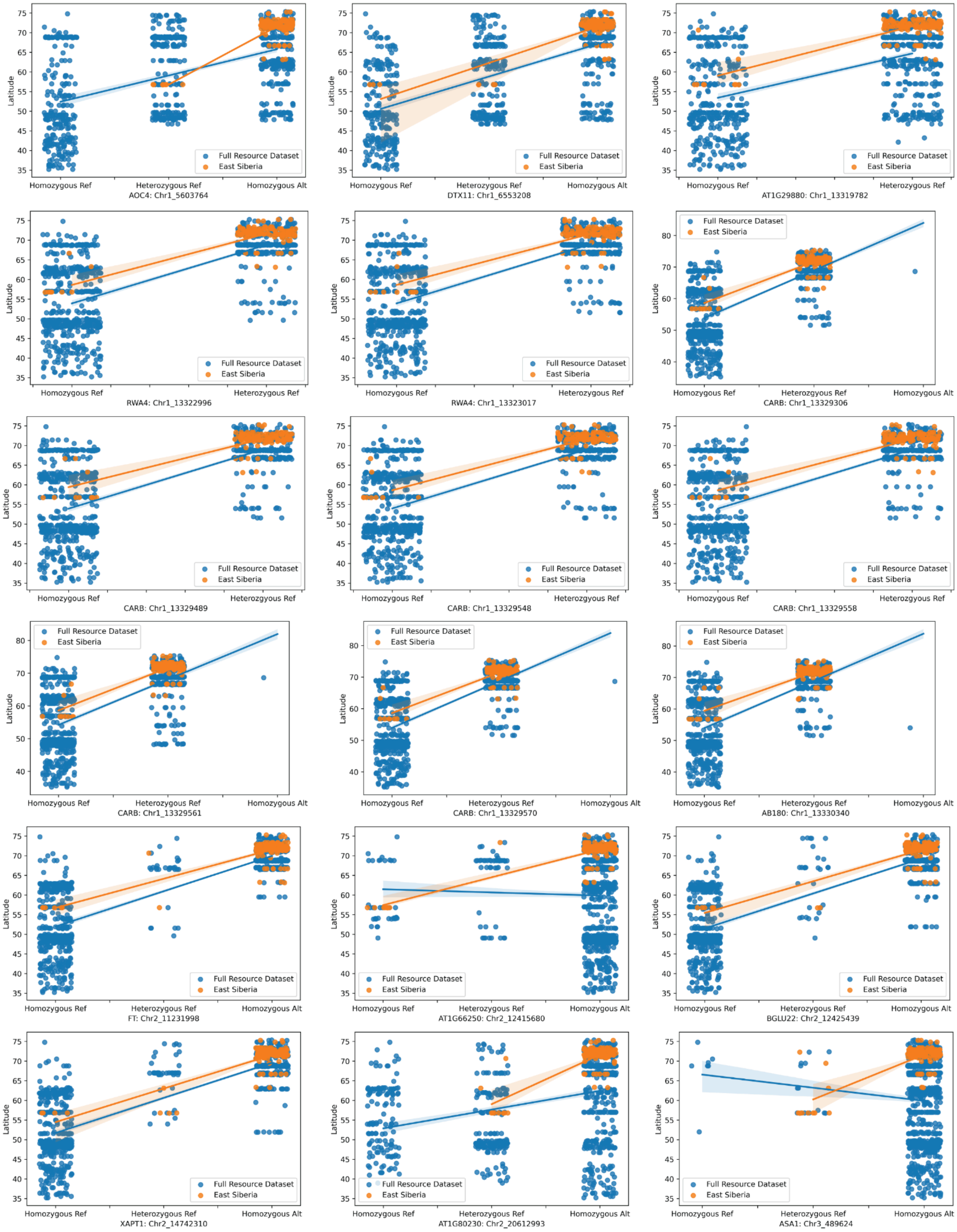
Latitude
Full Resource Dataset
East Siberia
Homozygous Ref
Heterozygous Ref
Homozygous Alt
AOC4: Chr1_5603764
DTX11: Chr1_6553208
AT1G29880: Chr1_13319782
RWA4: Chr1_13322996
RWA4: Chr1_13323017
CARB: Chr1_13329306
CARB: Chr1_13329489
CARB: Chr1_13329548
CARB: Chr1_13329558
CARB: Chr1_13329561
CARB: Chr1_13329570
AB180: Chr1_13330340
FT: Chr2_11231998
AT1G66250: Chr2_12415680
BGLU22: Chr2_12425439
XAPT1: Chr2_14742310
AT1G80230: Chr2_20612993
ASA1: Chr3_489624

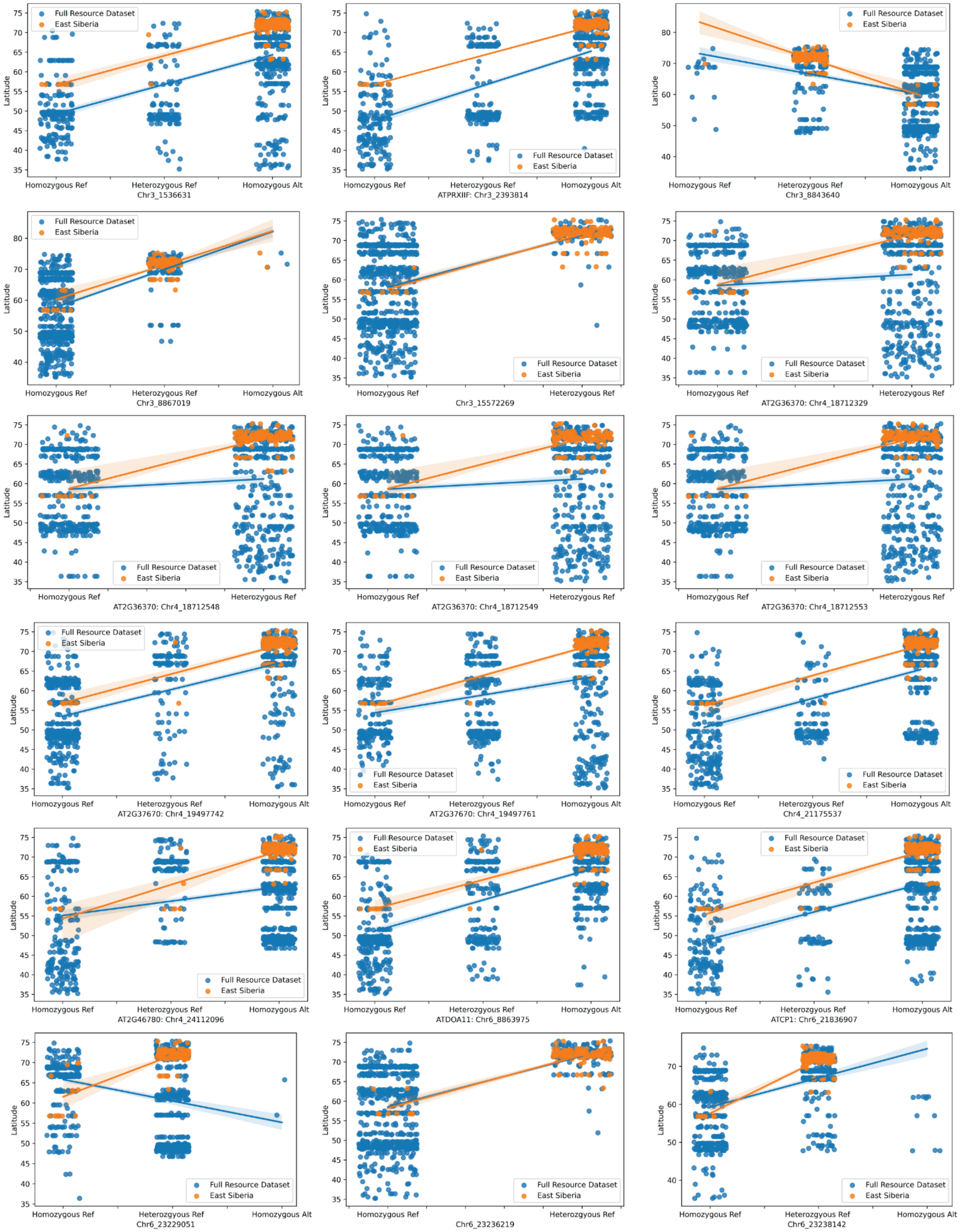
Full Resource Dataset
East Siberia
Latitude
Homozygous Ref
Heterozygous Ref
Homozygous Alt
Chr3_1536631
ATPRXIIF: Chr3_2393814
Chr3_8843640
Chr3_8867019
Chr3_15572269
AT2G36370: Chr4_18712329
AT2G36370: Chr4_18712548
AT2G36370: Chr4_18712549
AT2G36370: Chr4_18712553
AT2G37670: Chr4_19497742
AT2G37670: Chr4_19497761
Chr4_21175537
AT2G46780: Chr4_24112096
ATDOA11: Chr6_8863975
ATCP1: Chr6_21836907
Chr6_23229051
Chr6_23236219
Chr6_23238142

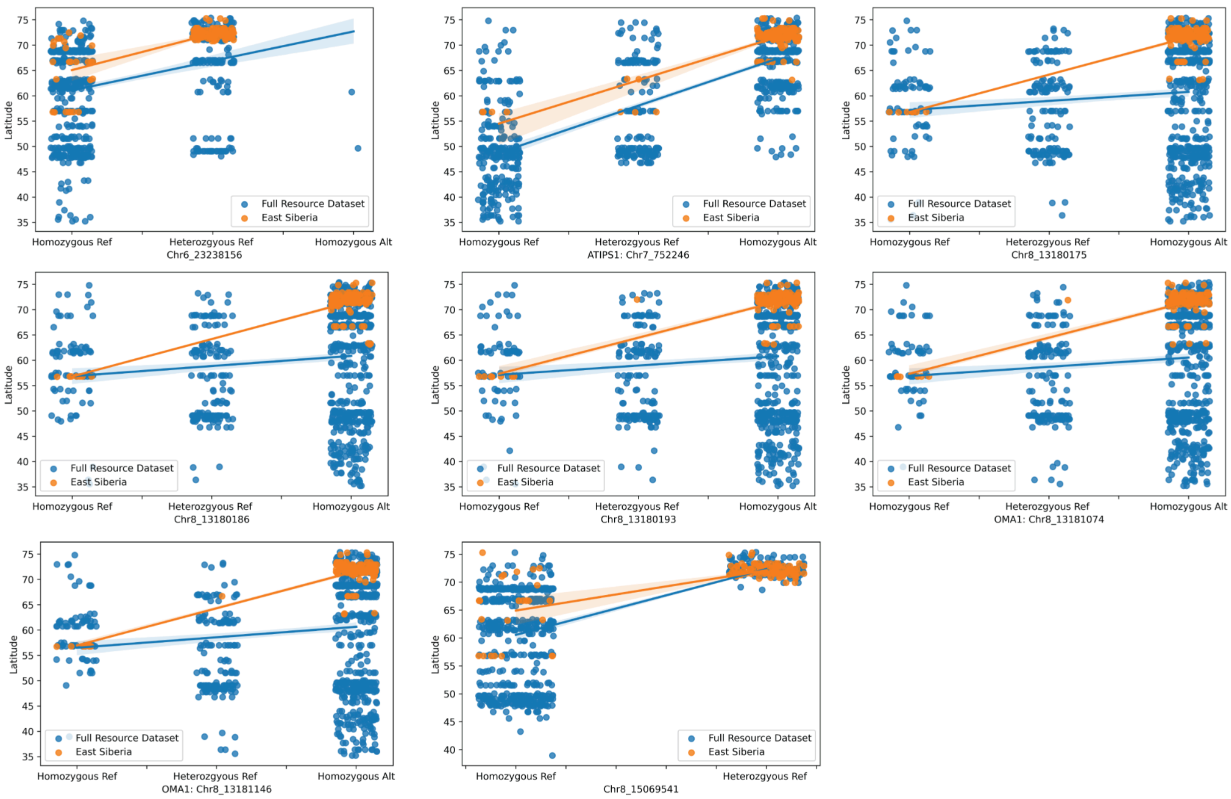


Supplementary fig. 7 Linear regression of latitude and genotypes of candidate genes within Eastern Siberian lineage (orange) and across the entire *A. lyrata* dataset (blue).

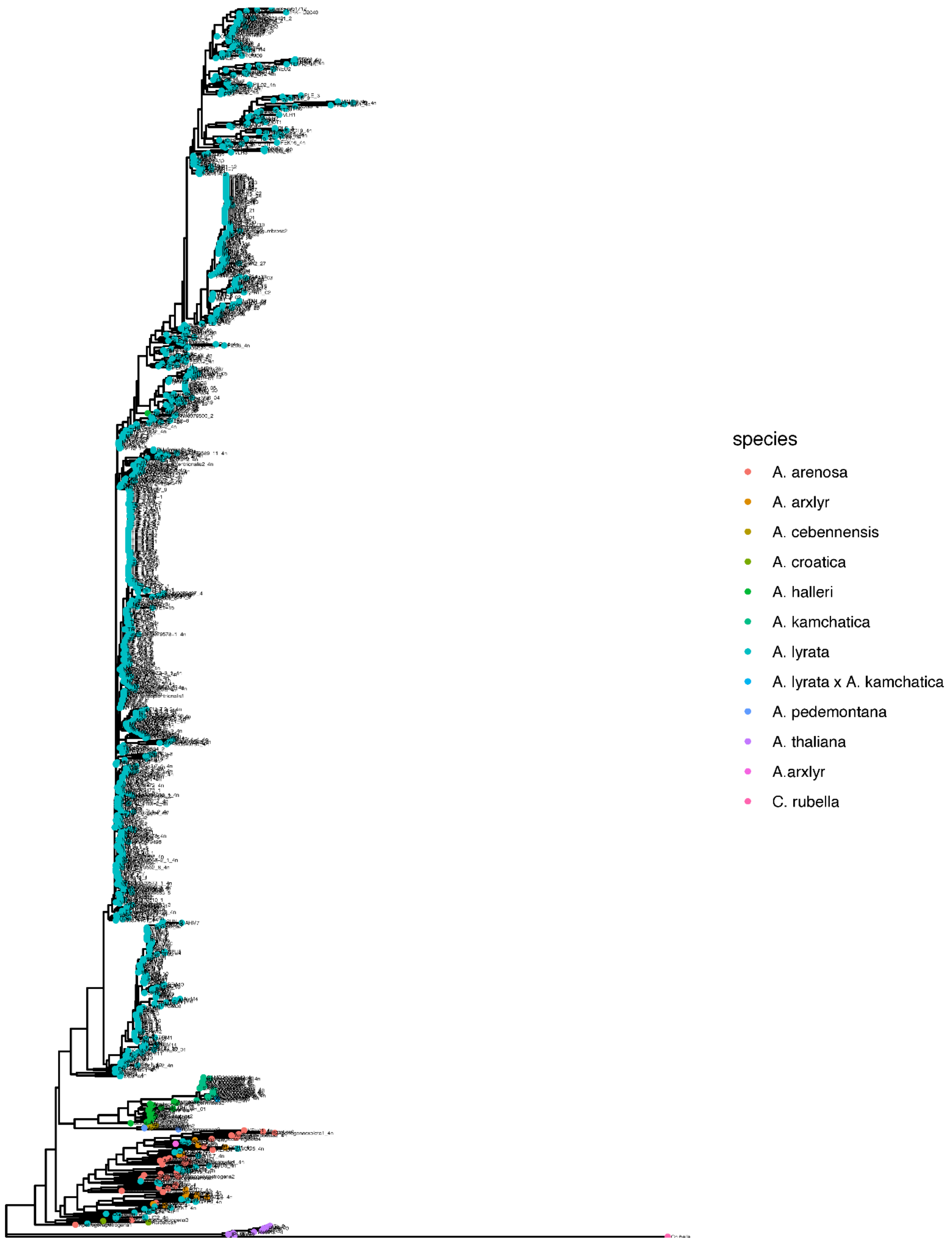


Supplementary fig. 8. Phylogenetic gene tree of ATIPS1 DNA sequences for the full *A. lyrata* dataset with outgroup taxa designated for SNP polarization relative to the *A. lyrata* reference allele. The top *A. lyrata*-specific cluster represents a derived and “Northern” allele, the bottom cluster is mixed and represents the ancestral “southern” allele.
